# tauFisher accurately predicts circadian time from a single sample of bulk and single-cell transcriptomic data

**DOI:** 10.1101/2023.04.04.535473

**Authors:** Junyan Duan, Michelle N. Ngo, Satya Swaroop Karri, Lam C. Tsoi, Johann E. Gudjonsson, Babak Shahbaba, John Lowengrub, Bogi Andersen

## Abstract

As the circadian clock regulates fundamental biological processes, disrupted clocks are often observed in patients and diseased tissues. Determining the circadian time of the patient or the tissue of focus is essential in circadian medicine and research. Here we present tauFisher, a computational pipeline that accurately predicts circadian time from a single transcriptomic sample by finding correlations between rhythmic genes within the sample. We demonstrate tauFisher’s outstanding performance in both bulk and single-cell transcriptomic data collected from multiple tissue types and experimental settings. Application of tauFisher at a cell-type level in a single-cell RNA-seq dataset collected from mouse dermal skin implies that greater circadian phase heterogeneity may explain the dampened rhythm of collective core clock gene expression in dermal immune cells compared to dermal fibroblasts. Given its robustness and generalizability across assay platforms, experimental setups, and tissue types, as well as its potential application in single-cell RNA-seq data analysis, tauFisher is a promising tool that facilitates circadian medicine and research.

## 1 Introduction

Organisms have evolved intrinsic circadian clocks that help them anticipate and adjust to environmental changes caused by the 24-hour rotation of the Earth [1, 2]. The mammalian circadian clock is a biochemical oscillator powered by transcription-translation loops consisting of a positive arm and a negative arm [1–3]. In the positive arm, BMAL1 and CLOCK promote the expression of clock-controlled genes, including the negative arm factors PER and CRY. PER and CRY inhibit the activating effect of BMAL1-CLOCK, leading to 24-hour oscillations.

In mammals, the suprachiasmatic nucleus (SCN) of the hypothalamus is the central pacemaker that coordinates and synchronizes circadian rhythms in peripheral tissues through neuronal and hormonal signals [4]. Besides signals from the SCN, environmental signals such as temperature [4], feeding [5, 6], and direct light [7] can selectively set peripheral clocks, sometimes causing asynchrony between the central and peripheral clocks. Epidemiological studies of shift workers and chronically jet-lagged individuals show correlations between circadian disruption and cardiovascular diseases [8], mental health disorders [9], metabolic diseases [10–12], as well as cancer in various organs [13, 14], including skin [15, 16], breast [17, 18], and prostate [19, 20].

The goal of the nascent field of circadian medicine is to consider circadian rhythm and its disruption in patient care. As the rhythm of a patient or diseased tissue is not necessarily synchronized with the external light-dark cycle, an important challenge in circadian medicine is to determine the internal circadian time of the patient or the tissue of focus. Such information can determine optimal time of treatment and identify conditions that might benefit from restoring circadian function [21, 22]. Current methods of circadian timing determination for a patient include the dim-light melatonin-onset assay [23], as well as circadian rhythm inference from body temperature [24], or cortisol levels in biofluids [25].

Additionally, determining the circadian time of a sample is important for research. With the explosion of bulk and single-cell transcriptomics datasets, there is growing effort to integrate and compare such datasets. As about 10% of the transcriptome has diurnal expression patterns, analyzing such datasets without their time stamps may lead to inconsistent observations that are dependent on the time of sample collection. Hence, there is a need to develop method that can determine the circadian time of such datasets.

Several groups have developed methods to infer circadian time of a sample (organism, organ, or tissue) based on transcriptomic data. CYCLOPS [26, 27] infers circadian phases by ordering the data collected from the entire periodic cycle. Several methods were also developed to predict circadian time from a single sample. ZeitZeiger [28] identifies useful features (genes) for prediction, scales the features over time, applies sparse principal component analysis, and predicts according to maximum likelihood estimation. BIO CLOCK [29] uses supervised deep neural networks with coupled sine and cosine output units. TimeSignatR [30] applies within-subject renormalization and an elastic net predictor, making it generalizable between transcriptomic data from different assay platforms. Recently, a Bayesian variational inference approach called Tempo [31] was designed to predict circadian phase in single cell transcriptomics and it quantifies estimation uncertainty.

Among the methods mentioned above, ZeitZeiger, BIO CLOCK and TimeSignatR are the methods that can predict circadian phase from a single sample of bulk transcriptomic data and are generalizable for different tissues. But, they have limitations. ZeitZeiger frequently runs into linear dependency issues, needs to be retrained before each prediction, and is not generalizable between transcriptomic platforms. BIO CLOCK does not require re-training for each prediction but is not time-efficient. TimeSignatR performs well if there are two test samples, but performance depends on the time interval between the two samples. When given only one test sample, TimeSignatR can infer a second sample, but it has very low accuracy.

Here, we present tauFisher, a pipeline that can accurately predict circadian time from a single transcriptomic data irrespective of the transcriptomic platform. tauFisher improves on previous methods in several ways: (1) it does not require the training data to be a complete time series; (2) the within-sample normalization step allows tauFisher to give an accurate prediction from just one sample; (3) since tauFisher only needs a few features to make accurate predictions, training and testing are computationally efficient; (4) tauFisher is platform agnostic and users only need to train the predictor once and can use the same predictor to make predictions for external datasets of the same tissue, regardless of the platform; and (5) unprecedentedly, tauFisher trained on bulk sequencing data is able to accurately predict the circadian time of single-cell RNA sequencing (scRNA-seq) data, and it can be used to investigate circadian phase heterogeneity in different cell types.

We collected a time series of scRNA-seq data from mouse dermis in this study and found that most of the rhythmic processes are metabolism-related in dermal fibroblasts, whereas almost all rhythmic processes are related to immune responses in dermal immune cells. Additionally, we found that the amplitude of the collective rhythm is dampened in dermal immune cells compared to dermal fibroblasts. Incorporating tauFisher with bootstrapping revealed that circadian phase heterogeneity contributes to the dampened collective rhythm as well as fewer rhythmic genes found in dermal immune cells.

## 2 Results

### 2.1 Overview of tauFisher

tauFisher is an assay platform-agnostic method that predicts circadian time from a single transcriptomic sample. The training part of the pipeline, which requires a time series of transcriptomic data, consists of five main steps: (1) identifying diurnal genes with a period length of 24 hours, (2) curve fitting using functional data analysis to fill in the missing time points and to decrease noise in the training data, (3) within-sample normalization by calculating and scaling the difference in expression for each pair of predictor genes, (4) linearly transforming the scaled differences using principal component analysis, and (5) fitting a multinomial regression on the first two principal components (Figure 1, Method Section 4.1).

**Fig. 1.**
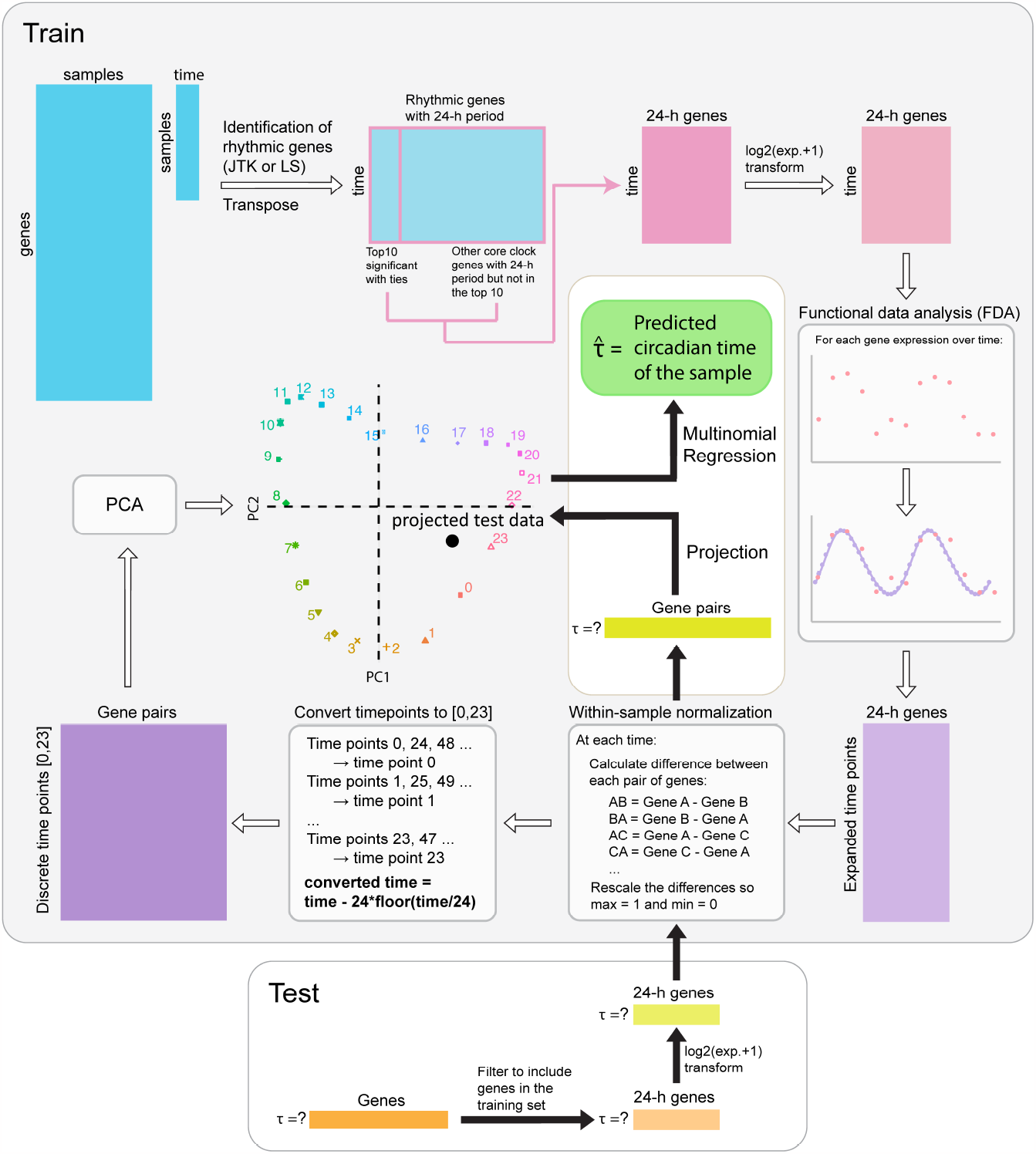
Key steps of the tauFisher pipeline includes identification of periodic genes, functional data analysis, within-sample feature normalization, linear transformation, and multinomial regression.

For testing, a transcriptomic sample without a time label is narrowed to include only the predictor genes identified in the training data. After the within-sample normalization step, the test sample is projected to the principal component space and multinomial regression is performed to predict time of the test sample (Figure 1, Method Section 4.1).

### 2.2 tauFisher outperforms current methods when trained and tested on bulk-sequencing data

To assess the robustness and accuracy of tauFisher in predicting circadian time from a single sample of transcriptomic data, we applied tauFisher to a diverse set of data collected from different species, tissues and sequencing platforms (Table 1).

**Table 1.** Datasets from different species, tissues and sequencing platforms were used to benchmark tauFisher’s ability to predict circadian time.

| Data | Year | GEO | Species | Tissue | Platform | Sampling Frequency | Time Course Duration |
| --- | --- | --- | --- | --- | --- | --- | --- |
| Arnardottir ES *et al.* [32] | 2014 | GSE56931 | *Homo sapiens* | Blood | Custom Affymetrix Microarray | 4h | 72h |
| Braun R *et al.* [30] | 2018 | GSE113883 | *Homo sapiens* | Blood | Illumina NextSeq 500 | 2h | 28h |
| Geyfman M *et al.* [33] | 2012 | GSE38622 | *Mus musculus* | Skin | Affymetrix Mouse Gene 1.0 ST Array | 4h | 48h |
| Tognini P *et al.* [34] | 2020 | GSE157077 | *Mus musculus* | SCN | Illumina HiSeq 4000 | 4h | 24h |
| Zhang R *et al.* [35] | 2014 | GSE54650 | *Mus musculus* | Kidney | Affymetrix Mouse Gene 1.0 ST Array | 2h | 48h |
| Zhang R *et al.* [35] | 2014 | GSE54650 | *Mus musculus* | Liver | Affymetrix Mouse Gene 1.0 ST Array | 2h | 48h |
| Zhang R *et al.* [35] | 2014 | GSE54651 | *Mus musculus* | Kidney | Illumina HiSeq 2000 | 6h | 48h |
| Zhang R *et al.* [35] | 2014 | GSE54651 | *Mus musculus* | Liver | Illumina HiSeq 2000 | 6h | 48h |

For each benchmark dataset, we generated 100 random train and test partitions (without replacement) of the samples. In each partition, we used 80% of the samples for training and 20% for testing.

We compared tauFisher to the current state-of-the-art methods: ZeitZeiger [28] and TimeSignatR [30]. As TimeSignatR is able to predict circadian time for samples from one time point alone, or one time point with an inferred second time point, we included both types of predictions in the benchmark.

We define a prediction within two hours of the true time to be correct. Using other time ranges to define correctness minimally change the benchmark outcome (Supplementary Table 1).

tauFisher achieved the highest accuracy for all eight benchmark datasets; seven using predictor genes found by JTK Cycle [36] and one using Lomb-Scargle [37] (Figure 2; Supplementary Table 2). While ZeitZeiger did not attain the highest accuracy in any of the datasets, it achieved the lowest root mean squared error (RMSE) in three of the datasets and comparable RMSE to tauFisher in half of the datasets. However, ZeitZeiger could not predict the time for several iterations due to linearly dependent basis vectors. Particularly, in the kidney and liver bulk sequencing datasets, ZeitZeiger failed to predict the time for all 100 iterations. TimeSignatR performed the worst among the three methods, giving the lowest accuracy and the highest RMSE.

**Fig. 2.**
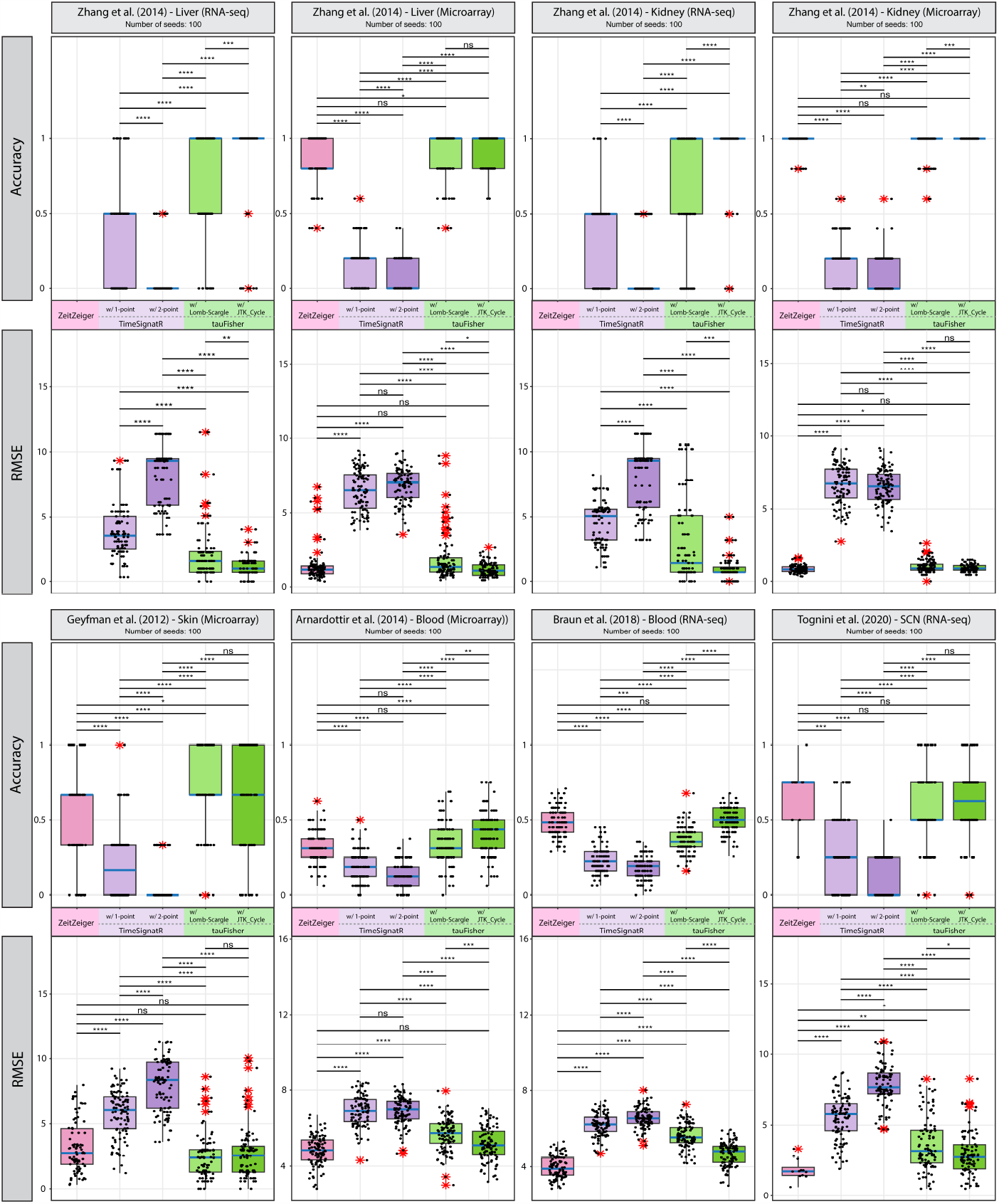
tauFisher outperforms published circadian time prediction methods in both accuracy and RMSE for transcriptomic data collected from various organs and assay platforms. ns: *p*-value *>* 0.05, ∗: *p*-value ≤ 0.05, *p*-value ≤ 0.01, ∗ ∗ ∗: *p*-value ≤ 0.001, ∗ ∗ ∗∗: *p*-value ≤0.0001.

### 2.3 tauFisher accurately predicts circadian time for cross-platform bulk transcriptomic data

Since tauFisher gives accurate circadian time prediction for bulk transcriptomic data collected from various platforms, we examined its performance when trained and tested on datasets generated from different platforms. We used rhythmic genes identified by JTK Cycle in the tauFisher pipeline, since this combination gave the most accurate predictions in the within-platform benchmark.

We trained tauFisher on GSE38622 [33], a microarray dataset collected from mouse dorsal skin every four hours for 48 hours under regular 12:12 light-dark cycle (zeitgeber time [ZT] 2, 6, 10, …). The test dataset is from GSE83855 [6], a bulk RNA-seq dataset collected every four hours for 28 hours under 12:12 light-dark cycle (ZT0, 4, 8, …) from mouse dorsal skin in a time-restricted feeding study. Since time of feeding influences tissue’s circadian clock [6, 22], we only selected the ad libidum control condition for testing so that the time labels best represent the internal time.

Eighteen genes were selected to be predictor features. Though the train and test datasets are not on the same scale and were collected at different time points, their overall rhythmic patterns agree with each other (Figure 3a). For seven out of the eight tests, tauFisher predicted a circadian time that is within the 2-hour range from the actual time label, giving a high accuracy of 0.875 and a low RMSE of 2.669 (Figure 3a). This example demonstrates tauFisher’s ability to accurately predict circadian time across bulk transcriptomics platforms.

**Fig. 3.**
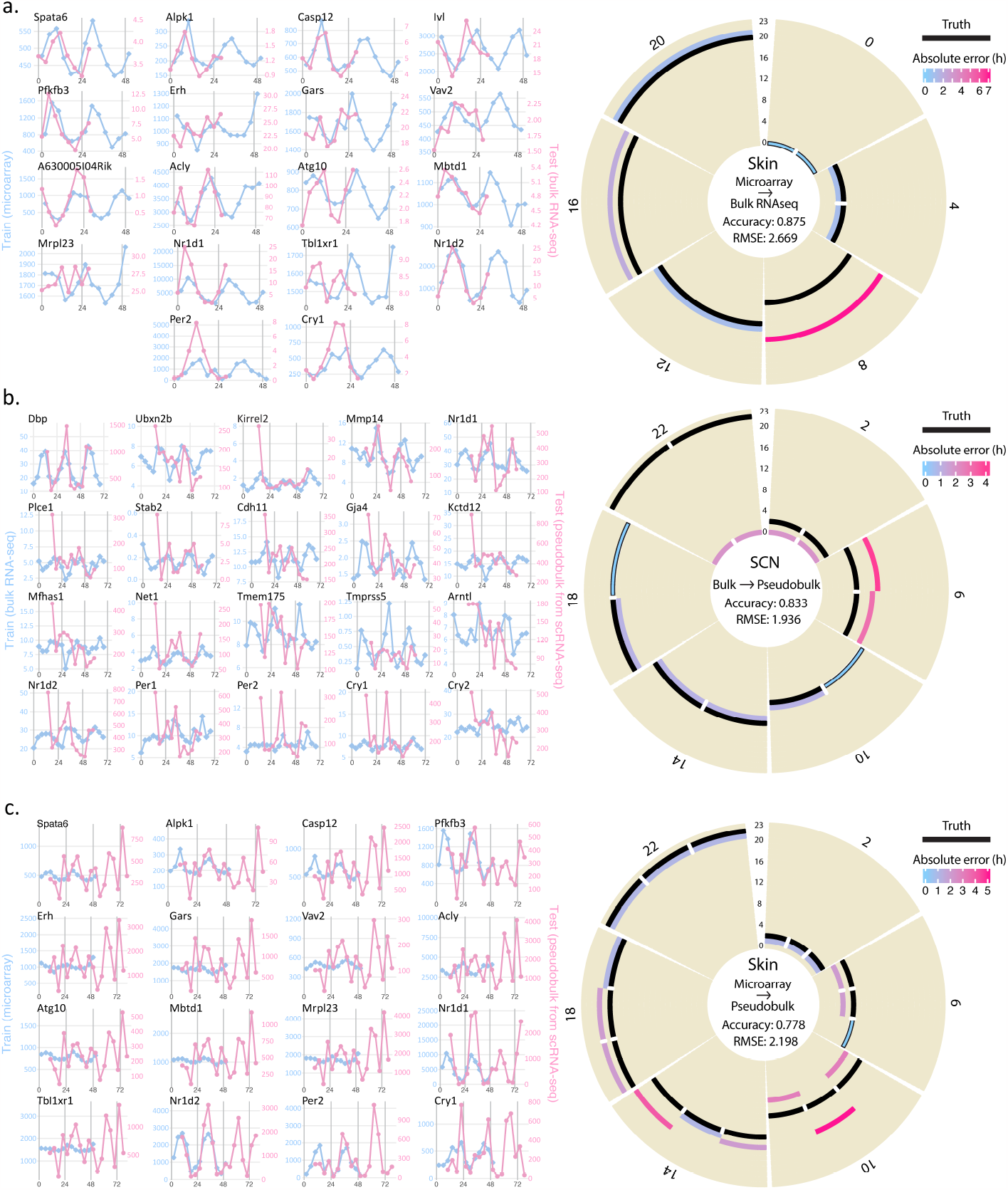
tauFisher accurately predicts circadian time even when the training and test data are from different assay methods. **a** tauFisher trained on skin microarray data can predict circadian time for skin bulk RNA-seq data. **b** tauFisher trained on SCN bulk RNA-seq data can predict circadian time of pseudobulk data generated from SCN scRNA-seq data. **c** tauFisher trained on skin microarray data can be used to predict circadian time of pseudobulk data generated from dermis scRNA-seq data. **a, b, c** Left: Predictor features and their expression in the training and test data. Right: Prediction outcomes for the tests. Labels on the circumference are the time labels for the test data. Predicted times are plotted along the radius, with colors representing absolute error. Black bars along the radius marks the truth (same as the labels on the circumference).

### 2.4 tauFisher trained on bulk RNA-seq data and microarray data accurately predicts circadian time of scRNA-seq samples

tauFisher’s ability to predict circadian time is not limited to cross-platform bulk-level transcriptomic datasets. It can predict circadian time for scRNA-seq data as well. In particular, tauFisher only needs to be trained on a time series of bulk-level transcriptomic data, which is more abundant and cheaper to collect than a scRNA-seq data time series.

Since most published scRNA-seq datasets do not have time labels, the selection of test datasets was limited. Here we tested tauFisher on scRNA-seq data collected from the mouse SCN [38] and mouse dermal skin (collected in this study).

GSE117295 [38] includes twelve single-cell SCN samples collected from circadian time (CT) 14 to 58 every four hours (CT14, 18, 22, …) under constant darkness, and one light-stimulated SCN sample. Since light immediately induces differential expression of rhythmic genes [38], only the samples from the control experiment were used for benchmarking. For each of the twelve samples, a pseudobulk dataset was generated for testing (Method Section 4.5). For training, we chose GSE157077 [34], a time series of bulk RNA-seq data collected from the mouse SCN every four hours under regular 12:12 light-dark cycle starting at ZT0. Since each time point in the training dataset contains three replicates, instead of averaging them, we concatenated the replicates so that the input training data spans 72 hours.

Twenty genes from the training data passed the feature selection criteria. These genes display robust rhythms in both the training data and the testing pseudobulk data (Figure 3b). The test data appears to be noisier since it is not normalized. tauFisher does not require the test data to be normalized as the within-sample normalization step is part of the pipeline.

In ten out of the twelve tests, tauFisher predicted a time that is within 2-hour of the labeled time, resulting a high accuracy of 0.833 and a low RMSE of 1.936 (Figure 3b). Although neither ZeitZeiger nor TimeSignatR claims to be able to predict time for scRNA-seq data, we still tested their performance. tauFisher greatly outperforms ZeitZeiger and TimeSignatR in both accuracy and RMSE (Table 2).

**Table 2.** tauFisher trained on SCN bulk RNA-seq data [34] accurately predicts circadian time for pseudobulk data (generated from scRNA-seq data [38]).

| Sample | Truth | ZeitZeiger |  | TimeSignatR<br>w/ 1pt |  | TimeSignatR<br>w/ 2pt |  | tauFisher<br>w/ JTK.Cycle |  |
| --- | --- | --- | --- | --- | --- | --- | --- | --- | --- |
|  |  | Prediction | Error | Prediction | Error | Prediction | Error | Prediction | Error |
| CT14 | 14 | 16 | -2 | 14 | 0 | 14 | 0 | 13 | 1 |
| CT18 | 18 | 2 | -8 | 12 | 6 | 15 | 3 | 17 | 1 |
| CT22 | 22 | 2 | -4 | 12 | 10 | 14 | 8 | 0 | -2 |
| CT26 | 2 | 2 | 0 | 11 | -9 | 13 | -11 | 0 | 2 |
| CT30 | 6 | 15 | -9 | 11 | -5 | 12 | -6 | 10 | -4 |
| CT34 | 10 | 15 | -5 | 11 | -1 | 12 | -2 | 10 | 0 |
| CT38 | 14 | 16 | -2 | 14 | 0 | 14 | 0 | 13 | 1 |
| CT42 | 18 | 2 | -8 | 13 | 5 | 15 | 3 | 18 | 0 |
| CT46 | 22 | 2 | -4 | 13 | 9 | 16 | 6 | 0 | -2 |
| CT50 | 2 | 2 | 0 | 11 | -9 | 16 | 10 | 0 | 2 |
| CT54 | 6 | 15 | -9 | 10 | -4 | 11 | -5 | 9 | -3 |
| CT58 | 10 | 15 | -5 | 12 | -2 | 12 | -2 | 11 | -1 |
| **Accuracy** |  | 0.33 |  | 0.33 |  | 0.33 |  | **0.83** |  |
| **RMSE** |  | 5.63 |  | 6.12 |  | 5.83 |  | **1.94** |  |

To ensure that tauFisher’s performance on scRNA-seq data is consistent and applicable to peripheral clocks, we performed scRNA-seq on adult wild type C57BL/6J mouse dorsal dermis every four hours for 72 hours under 12:12 light-dark cycle. The pseudobulk matrices for the 18 samples were computed directly from the unprocessed data. We trained tauFisher on GSE38622 [33], a time series of skin microarray data. Because two of the rhythmic genes, *A630005I04Rik* and *Ivl*, are not present in the pseudobulk data, only 16 features were selected in the tauFisher pipeline in this test (Figure 3c).

Although the input test data, the unnormalized pseudobulk data, appears to be noisy, tauFisher successfully predicts circadian times for the 18 samples thanks to the within-sample normalization step in the tauFisher pipeline. In 14 out of the 18 tests, tauFisher predicted circadian time within 2 hours of the labeled time, giving a high accuracy of 0.778 and a low RMSE of 2.198 (Figure 3c).

In sum, we have demonstrated that tauFisher trained on bulk-level transcriptomic data, either bulk RNA-seq or microarray data, can accurately predict circadian time for scRNA-seq data, making it particularly useful for expanding the current scRNA-seq database for circadian studies by adding time labels to existing scRNA-seq data.

### 2.5 Collective circadian rhythms are dampened in dermal immune cells compared to dermal fibroblasts

Due to the frequency of sequencing dropouts for clock genes in scRNA-seq data, investigating the circadian clock within each cell is not yet achievable. To overcome this limitation, previous studies have used pseudobulk approaches to investigate the clock in scRNA-seq data [38].

To validate the pseudobulk approach for studying the circadian clock in mouse dermis, we normalized the pseudobulk scRNA-seq data, and compared it with the published microarray data GSE38622 from mouse whole skin [33]. Overlay of the expression of nine core clock genes, *Arntl, Dbp, Per1, Per2, Per3, Nr1d1, Nr1d2, Cry1* and *Cry2*, reveals perfect consistency between the microarray data and the scRNA-seq data (Figure 4a), indicating that circadian clock gene expression in the dermis is captured in the pseudobulk data generated from scRNA-seq data.

**Fig. 4.**
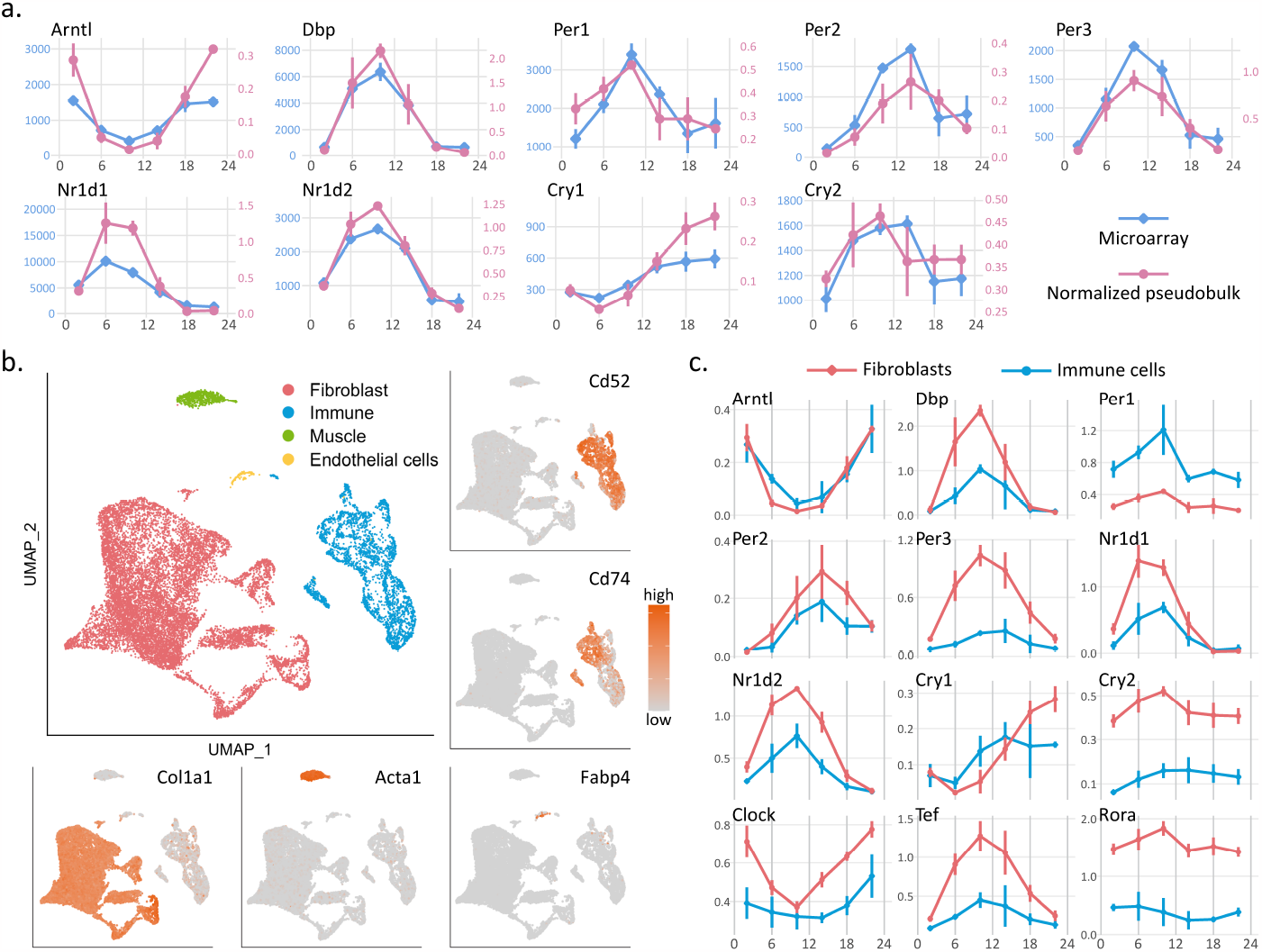
**a** Expression of the core clock genes in the normalized pseudobulk generated from scRNA-seq data (pink) is consistent with their expression in the published microarray data (blue). **b** Four major cell types, fibroblasts (red), immune cells (blue), muscle cells (green) and endothelial cells (yellow) are identified using canonical marker genes. Feature plots of the representative marker genes are shown (orange: high expression; grey: low expression); *Col1a1* for fibroblasts, *Acta1* for muscle cells, *Fabp4* for endothelial cells, *Cd52* and *Cd74* for immune cells. **c** At the pseudobulk-level, expression pattern of the core clock genes are similar in fibroblasts (red) and immune cells (blue), while the amplitudes of the oscillations are dampened in immune cells fore most of the core clock genes.

To study the circadian clock at a cell-type level in the skin, we integrated all samples and performed scRNA-seq analysis to identify cell types. In total, 16,866 cells passed the quality control, with around 950 cells per sample and around 2,800 cells per ZT. Four major cell types, fibroblasts (N = 12,649), immune cells (N = 3,353), muscle cells (N = 722) and endothelial cells (N = 142) were identified using canonical marker genes (Figure 4b). Due to low cell counts for muscle and endothelial cells (N*<*20) in some samples, we could not generate a reliable time series of pseudobulk data for these two cell types. Thus, we focus on the circadian clock in dermal fibroblasts and immune cells in this study.

To compare the core clock in fibroblasts and immune cells, we computed and normalized the pseudobulk data for each of the two cell types in each sample. Both fibroblasts and immune cells possess robust circadian clock at the pseudobulk level. While the overall rhythms in the two cells types are consistent with each other, with core clock gene expressions peaking and troughing around the same time, the amplitudes of the oscillations are reduced in the immune cells compared to fibroblasts, indicating a dampened collective clock in immune cells (Figure 4c). Whether this observation indicates less synchronous clocks in immune cells than in fibroblasts, or weaker clock function in each individual immune cell, is not known.

### 2.6 Dermal fibroblasts and immune cells harbor different rhythmic pathways and processes

To study diurnal genes and pathways in dermal fibroblasts and immune cells, we used JTK Cycle to identify rhythmic genes from the normalized pseudobulk data. We identified 1,946 and 432 rhythmic genes in fibroblasts (Supplementary Table 3) and immune cells (Supplementary Table 4), respectively (Figure 5a). The fewer rhythmic genes in immune cells is not caused by the lower cell count of immune cells, as randomly down-sampling the fibroblasts to the number of immune cells produced similar results. Only 79 genes were rhythmic in both cell types, with most of them related to the core clock network and metabolism.

**Fig. 5.**
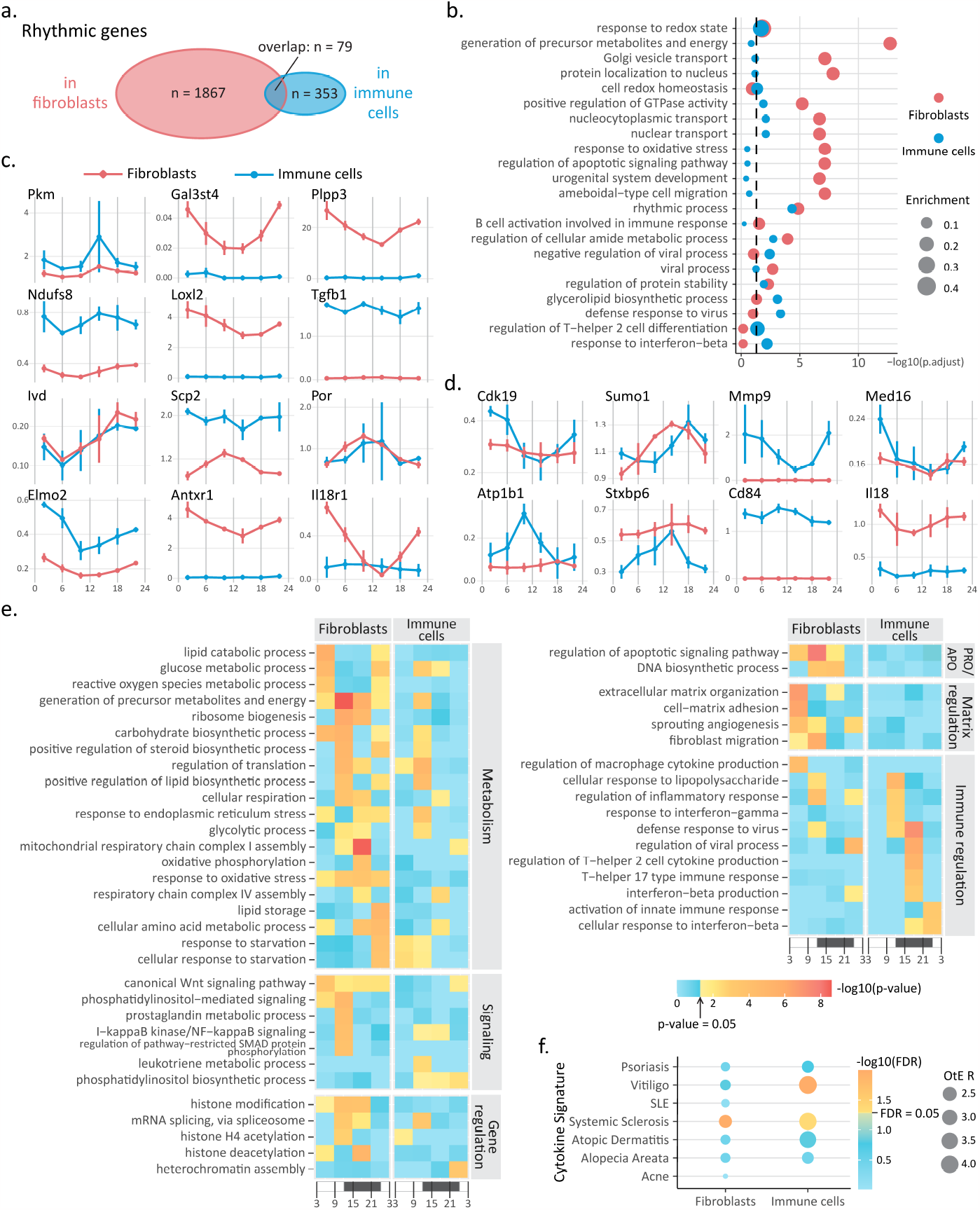
**a** JTK Cycle identified 1946 rhythmic genes in dermal fibroblasts (red) and 432 rhythmic genes in dermal immune cells (blue). Only 79 rhythmic genes are shared by the two cell types. **b** Gene Ontology analysis performed on rhythmic genes in fibroblasts (red) and immune cells (blue) reveals divergent biological processes being diurnally regulated in the two cell types. Dot size represents enrichment score. The vertical dashed line marks adjusted *p*-value = 0.05. **c, d** Expression of some of the rhythmic genes found in fibroblasts (**c**), and immune cells (**d**). **e** A heatmap showing *p*-values for some of the biological processes enriched by rhythmic genes peaking during each quarter-day time range. Color represents *p*-value. Blue: insignificant; yellow to red: significant with red representing lower *p*-value. x-axis represents time, with white being day and black being night. **f** Rhythmic genes in mouse dermal fibroblast and immune cells significantly enrich for genes within 200kb of the GWAS signals of immune-mediated skin conditions. FDR: false discovery rate; OtE R: observed-to-expected ratio. Blue: insignificant; yellow to orange: significant with orange representing lower FDR.

Gene Ontology analysis revealed that rhythmic processes in fibroblasts and immune cells are different. Shared terms reflect basic cell integrity maintenance and function, including *nucleocytoplasmic transport, regulations of cellular amide metabolic process, regulation of protein stability*, and *rhythmic process* (Figure 5b). For fibroblasts, additional metabolism processes and migration are significantly enriched in the rhythmic genes (Figure 5b, red). For immune cells, the rhythmic genes enrich for immune responses including *defense response to virus, regulation of T-helper 2 cell differentiation*, and *response to interferon-beta* (Figure 5b, blue).

We selected some of the rhythmic genes in fibroblasts (Figure 5c) and immune cells (Figure 5d) and compared their expression patterns in the two cell types. For fibroblasts, we highlight genes related to glucose metabolism (*Pkm*), glycosylation (*Gal3st4, Plpp3*), oxidative phosphorylation (*Ndufs8*), collagen regulations (*Loxl2, Tgfb1*), amino acid metabolism (*Ivd*), sterol synthesis (*Scp2, Por*), and cell adhesion and migration (*Elmo2, Antxr1*), suggesting circadian regulation of the above processes at a molecular level (Figure 5c,

Supplementary Table 3). Interestingly, while some genes are only significantly rhythmic in fibroblasts because they are not expressed in immune cells (e.g. *Loxl2*), some are expressed at similar or higher levels in immune cells, but are not significantly rhythmic in the latter (e.g. *Ndufs8, Scp2*), indicating cell-type specific circadian regulations.

In the immune cells, genes related to inflammatory and immune response (*Cdk19, Cd84*), post-translational modification (*Sumo1*), extracellular matrix regulation (*Mmp9*), transcription regulation (*Med16*), electrochemical gradient maintenance (*Atp1b1*), and intercellular communication (*Stxbp6*) are rhythmic (Figure 5d, Supplementary Table 4). We note that *Sumo1* is rhythmic in both fibroblasts and immune cells, but the expression peaks 4-hour later in immune cells than in fibroblasts.

Interestingly, the expression of *Il18r1* is significantly rhythmic with a large amplitude in fibroblasts (*p*-value = 2.21 × 10^−7^), but not in immune cells (*p*-value = 0.7104) (Figure 5c). The level of IL18, the ligand that binds to IL18R1, was found to be rhythmic in mouse peripheral blood [39]. Here, *Il18*, is significantly rhythmic in neither fibroblasts (*p*-value = 0.3097) nor immune cells (*p*-value = 0.0925) (Figure 5d). But, it is possible that the insignificance of the *p*-value for immune cells is caused by noise introduced by summing the expression of all types of immune cells while it is mostly expressed in the myeloid cells.

To further explore the rhythmic pathways in dermal fibroblasts and immune cells, we divided the list of rhythmic genes into four groups based on their peaking time (Method Section 4.5.3): day (ZT3 - ZT9), evening (ZT9 - ZT15), night (ZT15 - ZT21), and morning (ZT21 - ZT3 of the next day). The rhythmic genes are roughly evenly split: in fibroblasts, 426 peak during the day, 554 peak in the evening, 545 peak at night, and 421 peak in the morning (Supplementary Table 3); in immune cells, 129 peak during the day, 111 peak in the evening, 87 peak at night, and 105 peak in the morning (Supplementary Table 4). We then performed Gene Ontology analysis on the quarter-day rhythmic gene lists to identify the biological processes that are upregulated at different times of the day. We highlight some of the terms related to metabolism, signaling, cell proliferation and apoptosis, gene regulation, and immune regulation (Figure 5e).

During evening and night, when mice wake up, start feeding, and become active, processes such as *generation of precursor metabolites and energy, cellular respiration, mitochondrial respiratory chain complex I assembly* are upregulated in fibroblasts (Supplementary Table 5). Meanwhile, *glycolytic processes* are upregulated in fibroblasts, which is consistent with the finding that glycolysis is preferred at night in epidermal stem cells [40]. Additionally, similar to epidermal stem cells, more dermal fibroblasts may be in the S-phase of the cell cycle during the evening and night, as *DNA biosynthetic process* is enriched during this time. Various signaling pathways are also enriched during this time, including *prostaglandin metabolic process* and *regulation of apoptotic signaling pathway*. Gene-regulatory mechanisms such as *histone modification* and *mRNA splicing* are upregulated during the evening and night in fibroblasts. *Fibroblast migration* peaks at night, which is consistent with previous findings that mouse wounds heal fastest during the active phase [41]. Immune regulation is also circadian regulated in fibroblasts, as terms including *regulation of inflammatory response* are enriched during this time. Compared to dermal fibroblasts during the evening and night, fewer terms related to metabolism, signaling, and gene regulation are enriched in dermal immune cells (Supplementary Table 6). But, almost all immune regulation terms such as *defense response to virus* and *interferon-beta production* are upregulated in dermal immune cells during the evening and night, potentially contributing to shorter healing duration for wounds occurring during mice’s active phase as well [41]. Additionally, such findings in mice imply that circadian regulation of immune response may be related to the more severe symptoms of inflammatory skin diseases, such as psoriasis, in the evening and at night [22, 42].

In the morning and during the day, mice sleep and have lower food intake. Consistently, rhythmic genes peaking during this time in fibroblasts enrich for *lipid catabolic process, glucose metabolic process, lipid storage*, and *response to starvation*. Interestingly, *extracellular matrix organization* and *cell-matrix adhesion* peak during the day, possibly preparing for *fibroblast migration* which peaks in the evening (Supplementary Table 5). For immune cells, rhythmic genes peaking during the morning and day generate fewer terms than the ones peaking during the evening and night, especially in the immune regulation category (Supplementary Table 6). Interestingly, rhythmic genes the mouse dermal fibroblasts significantly enrich for genes linked to SNPs associated with systemic sclerosis [43], an inflammatory disease with increased collagen production by fibroblasts. Rhythmic genes in mouse dermal immune cells significantly enrich for SNPs associated with not only systemic sclerosis, but also vitiligo [44], a disease characterized by immune-mediated depigmentation of the skin (Figure 5f).

In sum, we found that more genes are collectively rhythmic in fibroblasts than in immune cells, while only a few rhythmic genes are shared. Additionally, more metabolism processes are diurnally regulated in fibroblasts, with respiration peaking during the evening and night, and response to starvation and lipid storage peaking during the morning and day. On other hand, immune regulation is almost exclusively upregulated by rhythmic genes that peak during the evening and night in immune cells. Importantly, rhythmic genes found in both fibroblasts and immune cells significantly enrich for Genomewide Association Studies (GWAS) SNPs associated with human skin immune-mediated conditions, pointing to a potential link between the skin circadian clock and autoimmune diseases of the skin.

### 2.7 tauFisher determines that circadian phases are more heterogeneous in dermal immune cells than in fibroblasts

Analysis of the pseudobulk data from dermal fibroblasts and immune cells reveals dampened amplitude of core clock genes (Figure 4d), and finds fewer rhythmic genes in immune cells than in fibroblasts (Figure 5a). This could mean that each individual immune cell harbors weaker circadian clock, and/or the immune cells have more heterogeneous phases, so collectively they display a dampened clock.

To investigate the cause behind the dampened clock in dermal immune cells, we executed a bootstrapping approach that incorporates tauFisher for its ability to predict circadian time for transcriptomic data at different scales (Figure 6a). Since the heterogeneity of a set of heterogeneous clocks should be captured at any given time point, the pipeline compares circadian phase heterogeneity in different cell types within each time point. At each time point, we select the cell types to be examined in the scRNA-seq data and limit the expression matrix so that it only contains the predictor genes identified in the training data. To remove influence from the difference in the number of cells, we randomly sample the same number of cells for each cell type, summing the transcript counts for each gene to create a pseudobulk dataset. We repeat the random sampling process to create pseudobulk replicates for each cell type. tauFisher then predicts circadian time for the pseudobulk replicates. Since the prediction outcomes are circular data, we then perform Rao’s Tests for Homogeneity to compare the mean and Wallraff Test of Angular Distances to compare the dispersion around the mean.

**Fig. 6.**
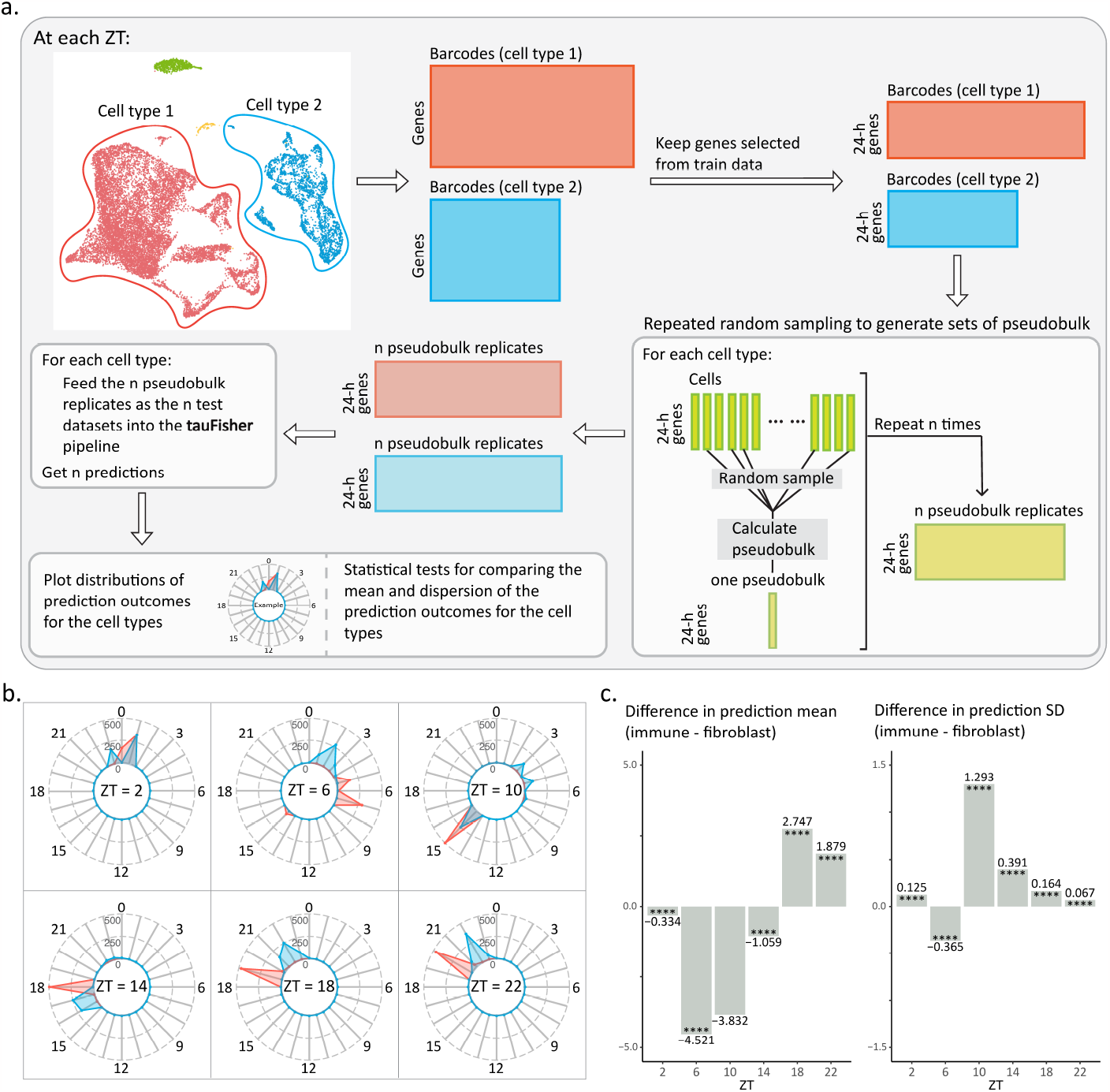
**a** A general overview of the bootstrapping approach we took to gain insight into circadian heterogeneity in phase in different cell types. **b** Radar plots showing the distribution of the prediction outcome for 500 pseudobulk replicates from dermal fibroblasts (red) and immune cells (blue). **c** Bar plots showing the differences between the prediction mean (left) and standard deviation (right) at each ZT (immune cells - fibroblasts). ∗ ∗ ∗∗: *p*-value ≤ 0.0001

To ensure that the pipeline works as expected, we generated simulated single-cell circadian gene expression datasets to represent a group of synchronized but dampened clocks, and a group of out-of-phase but robust clocks (Supplementary Figure 1). As expected, the prediction outcome for the out-of-phase clocks has a significantly greater dispersion around the mean, indicating a more heterogeneous mixture of phases (Supplementary Figure 1). We then perform the pipeline on the scRNA-seq data we collected, focusing on the fibroblasts and immune cells. At each time point, we randomly selected *n* cells for each cell type, with *n* equals to 20% of the cell count of the cell type with the minimum cell count. Then, we generated 500 pseudobulk replicates for each cell type at each time point, and used tauFisher to predict the circadian time for each replicate. We then compared the distribution of the prediction outcomes from the two cell types at each time point. In general, the prediction means are centered at different times for fibroblasts and immune cells (Figure 6b), but around the predicted time for the whole-sample pseudobulk data (Figure 3c). Whether one cell type’s circadian clock is ahead of the other is inconclusive (Figure 6c). Additionally, the distributions of the prediction outcome for immune cells are mostly multimodal and not as centered as the prediction distribution for fibroblasts (Figure 6b). Indeed, the standard deviation in the prediction distribution is significantly greater for five out of the six ZTs, indicating that immune cells have a more heterogeneous clock phase than fibroblasts.

In sum, we were able to use tauFisher to obtain insights into the circadian heterogeneity for different cell types by predicting the circadian time for random samples from each of the cell types. We hypothesize that the circadian clock is more heterogeneous in dermal immune cells than in dermal fibroblasts, and such heterogeneity may be the reason behind the dampened core clock and fewer rhythmic genes we found in immune cells based on collective, cell-type level, gene expression data. Such a result is not unexpected, as the fibroblasts may be more homogeneous in their biological function than the immune cells, which contain dendritic cells as well as different types of macrophages and lymphocytes (Supplementary Figure 2) that serve different immune functions. Unfortunately, we did not capture enough cells for each specific immune cell types in the scRNA-seq experiment to generate reliable pseudobulk data that is required for further circadian analysis (Supplementary Figure 2).

## 3 Discussion

In this study, we developed tauFisher, a computational pipeline that accurately predicts circadian time from a single transcriptomic dataset applicable to within-platform and cross-platform training-testing scenarios. Particularly, tauFisher trained on bulk transcriptomic data accurately predicts time for scRNAseq datasets. This method allows investigators to place a time stamp on transcriptomic datasets and facilitates the determination of circadian time in the context of circadian medicine.

Most transcriptomic datasets in public genomics repositories lack circadian time labels, which complicates integration with or comparison to other datasets. Adding time labels for existing transcriptomic datasets is important, as the clock modulates the expression of many protein-coding genes; it is necessary to know whether a significant gene is truly differentially regulated by a condition or the expression appears to be different because the samples were collected at different times. Also, computationally adding circadian time stamps to existing transcriptomic datasets collected from various platforms, including from scRNA-seq, opens up new possibilities for circadian research and allows investigators to take the full advantage of the shared resource in an efficient and inexpensive way.

Circadian time determination is also a key step in the implementation of circadian medicine. To maximize effectiveness while minimizing side effects of treatments, it is necessary to take into consideration the patient’s and relevant tissue’s actual circadian time. For example, on-pump cardiac surgeries in the afternoon are less likely to cause perioperative myocardial injury than when conducted in the morning [45] and cancer radiation therapy in the morning causes less skin damage than in the afternoon [46].

There have been several predictors of circadian time based on transcriptomic data [28–30], but to ensure wide applicability of this approach, a sequencing platform-agnostic method requiring low number of test samples is desired. tauFisher is preferable as it can accurately determine circadian time from a single sample of transcriptomic data collected from various assay methods including scRNAseq.

Once trained, tauFisher requires a single transcriptomic sample to predict the circadian time. We examined tauFisher’s ability to predict circadian time when the training and test data are from the same study, and we benchmarked it against state-of-the-art methods ZeitZeiger [28] and TimeSignatR [30]. tauFisher performed the best in accuracy and RMSE for almost all datasets. ZeitZeiger failed to run for several of the datasets due to linear dependency issues. When it did run, ZeitZeiger achieves similar accuracy and RMSE as tauFisher, possibly because both predictors build on principal component analysis, suggesting that the molecular clock is well captured and represented by orthogonal linear combinations of feature genes. TimeSignatR performed less well in the benchmark, even when a second sample was inferred. For time inference from a single sample, we reported worse general performance for TimeSignatR than reported in [30]. This could be due to the difference in the training data. In [30], TimeSignatR was trained on an independent dataset (GSE39445) from a different study [47] before testing on GSE56931 and GSE113883. Here we trained and tested TimeSingatR within GSE56931 and GSE113883. Authors of TimeSignatR also reported more accurate predictions by TimeSignatR with 2-point calibration than with just 1-point [30]. In this study’s benchmarking, TimeSignatR with 2-point calibration performed worse. Such discrepancy in performance reported between this study and [30] could be due to our implementation of the two-point calibration step in the TimeSignatR. In their vignette, the authors use the entire dataset (train and test data) during this two-point calibration step instead of just the training set. As tauFisher and ZeitZeiger do not utilize the test data in the processing of the train data, to make a fair comparison of their performance in predicting circadian time from one single test sample, we do not make the test sample available to the predictor until the actual testing step when implementing all three methods.

One of the most powerful features of tauFisher is its ability to accurately predict circadian time when trained and tested on datasets collected from different assay platforms under different experimental settings. tauFisher achieves high accuracy and low RMSE in not only bulk-to-bulk cross-platform predictions, but also bulk-to-scRNA-seq predictions. The consistency in performance despite drastically different assay methods and experimental setups suggests that tauFisher captures and extracts the underlying biological correlations in gene expressions while minimizing the effects of the noise and variability introduced by subjects and technology.

Two key steps in tauFisher help achieve this: functional data analysis for training data and within-sample normalization for both training and test data. Functional data analysis for the training data enables tauFisher to remove minor noise, smooth the time expression curves, and generate the expression data between the sampled time points. The within-sample normalization step for both training and test data calculates the difference between each pair of predictor genes at a given time point so that the feature matrix is expanded while some baseline noise is removed. The differences between the genes are then re-scaled to be between 0 and 1 so that the data become unit-less. Doing so in parallel on the training and test datasets brings them individually to the same scale, instead of batch-correcting the train to the scale of the test or the opposite. This allows testing of independent datasets without re-training. We note that this within-sample normalization is different from the within-subject normalization in [30], which is based on mean expression calculated from multiple samples collected from more than one time point over the circadian cycle.

In addition to testing tauFisher on published datasets, we also collected a time series of scRNA-seq from mouse dermis. Consistent with previous findings [22, 48], the circadian rhythm is robustly present in the dermis and the oscillatory patterns of the core clock genes agree with published data [33]. Comparing the rhythmic genes in fibroblasts and immune cells, we found that only a few rhythmic genes are shared by the two cell types and many pathways and processes are rhythmically regulated in a cell type-specific manner. Shared diurnally regulated terms includes basic cellular functions and the rhythmic genes in fibroblasts have greater enrichment for metabolism related terms, whereas the rhythmic genes in immune cells have greater enrichment for immune responses.

Combining tauFisher with other methods can guide the application of circadian medicine by providing additional insights and explanation of clinically observed circadian dysfunction. Dampened clock gene expression has been observed in psoriasis-affected skin [49, 50], as well as in various types of cancer [51–56]. There is also evidence that restoring dampened circadian oscillations in diseased tissues can be effective in reducing cancer cell proliferation and tumor growth [54, 56].

There are two possible behind-the-scene causes of dampened circadian rhythms at a bulk level: First, the circadian rhythm is dampened in every cell, but the cells are synchronous to each other. Second, the clock is normally functioning in every cell, but the cells are out of phase relative to each other. Understanding which of the two scenarios is responsible for a dampened bulk-level clock gene expression is particularly important because in one case, it would be optimal to stimulate the clock to restore the circadian clock in the diseased tissue, while in the other case, synchronizing the clock is more suitable.

Here, we observed that the collective circadian rhythm in dermal immune cells is dampened compared to fibroblasts. We incorporate tauFisher with bootstrapping to investigate the cause behind the dampened collective circadian rhythm in dermal immune cells. tauFisher’s prediction outcome suggests that the circadian phases in dermal immune cells are more heterogeneous than those in dermal fibroblasts, and this heterogeneity may contribute to the dampened rhythm in immune cells at a collective level.

The advantages tauFisher brings goes beyond accurately adding time stamps when incorporated with other methods. For example, combining tauFisher with a batch-effect correction method may facilitate a cleaner integration and help minimize the effect of the circadian clock in transcriptomic data analysis. This approach harbors great potential as many efforts are going into integrating datasets from different studies to create meta databases such as in the Human Cell Atlas.

In summary, tauFisher’s consistent and robust performance in accurately predicting circadian time from a single transcriptomic data makes it a useful addition to the toolbox of circadian medicine and research.

## 4 Methods

### 4.1 tauFisher

tauFisher is a platform-agnostic method that predicts the circadian time from a single transcriptomic sample. The method consists of three main steps: (1) identifying a subset of diurnal genes with a period length of 24 hours, (2) calculating the difference in expression for each pair of genes, and (3) linearly transforming the differences using PCA and fitting a multinomial logistic regression on the first two principal components.

#### 4.1.1 Averaging the expression matrix

The first step in tauFisher is to average each transcriptome by its genes such that the averaged gene expression matrix consists only of unique genes. The subsequent training data should consist of a gene expression matrix *X* ∈ ℝ^*N ×P*^ with *N* unique genes and *P* samples with known time and a vector *τ* ∈ R^*P*^ of the corresponding time for each sample. For scRNA-seq data, this averaged transcriptome over all the cells will be referred to as pseudobulk data.

#### 4.1.2 Identifying the periodic genes

tauFisher specifies either the Lomb-Scargle [37] or JTK Cycle [36] method in the `meta2d` function from R package MetaCycle [57] to determine the periodic genes and then selects the top ten statistically significant genes with a 24-hour period. These periodic genes are then combined with the core clock genes that also have a period of 24 hours to create the set of predictor genes *M*. The core clock genes for consideration in tauFisher are: *Bmal1, Dbp, Nr1d1, Nr1d2, Per1, Per2, Per3, Cry1*, and *Cry2*.

#### 4.1.3 Subset and transform the data

Subsetting the averaged expression matrix *X* on the set of predictor genes M yields averaged gene expression matrix *X*^*′*^ ∈ ℝ^*M ×P*^ with *M* periodic genes and *P* samples with known time; the vector *τ* should stay the same. The matrix *X*^*′*^ is then log-transformed element-wise: *X*^*′*^ = log_2_(*X*^*′*^ + 1).

#### 4.1.4 Run Functional Data Analysis

Since experiments have different sampling intervals throughout a circadian cycle, tauFisher uses functional data analysis (FDA) to represent the discrete time points as continuous functions. This allows tauFisher to evaluate and predict the circadian time of the new samples at any time point and reduces the noise from the training samples.

Briefly, each gene *m* has a log-transformed measurement at discrete time points *t*_1_, …, *t*_*P*_ ∈ *τ* that are generally equally spaced but may not be. These discrete values are converted to a function *Z*_*m*_ with values *Z*_*m*_(*t*) for any time *t* using a Fourier basis expansion:

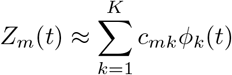

where *ϕ*_*k*_(*t*) is the *k*-th basis function for *k* = 1, …, *K* and ∀*t* ∈ *τ*, and *c*_*mk*_ is the corresponding coefficient. The Fourier basis is defined by *ϕ*_0_(*t*) = 1, *ϕ*_2*r*−1_(*t*) = sin(*rωt*), and *ϕ*_2*r*_(*t*) = cos(*rωt*) with the parameter *ω* determining the period 2*π/ω*. Since the log-transformed data matrix *X*^*I*^ is non-negative, a positive constraint is imposed such that the positive smoothing function is defined as the exponential of an unconstrained function: 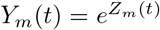. The smoothing function also contains a roughness penalty to prevent overfitting. In practice, tauFisher sets the number of basis functions to *K* = 5 because that produces curves that are the most sinusoidal however users can specify a different number of basis functions if they wish.

Although FDA represents the discrete time points as continuous functions for each gene, tauFisher predicts circadian time at user-defined time units instead of continuous time between 0 to 24. By default, the time units are set to discrete hours. The fitted functions *Y*_*m*_(*t*) are evaluated at the user-defined time units to create the smoothed expression matrix *Y*∈ ℝ^*×T*^, where *T* is the number of evaluated time points, and the new set of time points *τ*_*F*_ ∈ ℝ^*T*^. If the time course duration of the samples spans less than 24 hours, then the fitted curves are evaluated hourly from [0, 23] such that *T* = 24 to ensure all 24 hours are evaluated. If the time course duration of the samples spans greater than 24 hours, then fitted curves are evaluated from [*min*(*τ*), *max*(*τ*)] such that *T* = *max*(*τ*) − *min*(*τ*) + 1.

#### 4.1.5 Calculate differences between each gene pair

tauFisher then calculates the differences in the smoothed expression matrix *Y* for each pair of genes across time. Pairs of the same gene (i.e., the differences between Gene *a* and Gene *a*) are removed from the data matrix. Furthermore, tauFisher assumes that the effect of the difference between Gene *a* and Gene *b* and the effect of the difference between Gene *b* and Gene *a* on the predictor are not equal; thus, both pairs are included as covariates. For each time point *t*_1_, …, *t*_*T*_ ∈ *τ*_*F*_, these differences are then scaled to be [0, 1].

#### 4.1.6 Regress on the principal components

The differences matrix is projected onto a lower dimensional space via principal components analysis (PCA), and the first two principal components become covariates *x*_*i*1_ and *x*_*i*2_ for observation *i* in the multinomial regressor:

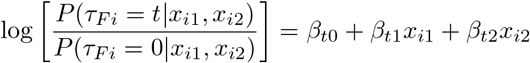

All time points *t*_1_, …, *t*_*T*_ ∈ *τ*_*F*_ are converted to be [0, 23], since time 0 is equal to time 24. Time zero, *τ*_*F*_ = 0, is set as the reference level in the model. The fitted multinomial regression model is then used to predict the circadian time of the new samples. We note that since time can be ordinal (accounting for the order) or continuous between [0, 24), we also tried other models such as an ordinal regression. However, these models were not as robust as the multinomial regression model and failed to run on the entire set of time points.

### 4.2 Calculating the difference in predicted time and true time

To evaluate the performance of tauFisher, we need to calculate how close the predicted time is to the true time. Since the outcome is cyclic and ranges from [0, 23], we apply the following conversion to calculate the true difference *D* from the difference *d* between the predicted time and true time:

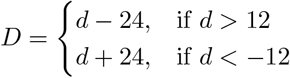

### 4.3 Simulating scRNA-seq circadian gene expression datasets

In Section 2.7, we verify that our pipeline can be used to investigate heterogeneous circadian phases using simulated scRNA-seq circadian gene expression data. We simulate two groups of data to represent two possible reasons for dampened expression: (1) a group of synchronized but dampened clock genes and (2) a group of normal (robust) but asynchronous clock genes. For both groups, the expressions of 9 representative “core clock” genes over a time course of 24-h are simulated using the following sine function:

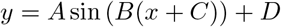

where *A* is the amplitude, *C* is the phase shift, *D* is the vertical shift, the period is 2*π/B*, and *x* is a sequence of integers from 0 to 23. We set *B* to be 24, such that the period is 2*π/*24, and *D* to be 25 to ensure that there are no negative gene expression values.

Previously, in Section 2.2, we used JTK Cycle [36] to identify the periodic genes of several datasets to benchmark tauFisher; part of JTK Cycle’s output is a data frame containing inferred amplitudes and phase shifts for each gene. So, as inputs for our simulated datasets, we select the inferred amplitude and phase shift values for core clock genes *Bmal1, Dbp, Nr1d1, Nr1d2, Per1, Per2, Per3, Cry1*, and *Cry2* from [34]. Then, for each dataset in Group (1), we simulate the expression of gene *i* as follows:

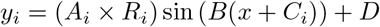

where *R*_*i*_ is one draw from a Beta(1, 2) distribution and all other parameters are as previously stated. Similarly, for each dataset in Group (2), we simulate simulate the expression of gene *i* as follows:

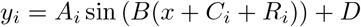

where *R*_*i*_ is one draw from a Normal(0, 12) distribution and all other parameters are also as previously stated. We generated 100 datasets for each group, which can be thought of as the simulated expression of 9 genes for 100 single cells over 24-h.

Since tauFisher is currently used for pseudobulk expression, we then convert our simulated single-cell datasets to two groups of 500 pseudobulk datasets (one for each case). We randomly select 6 time points without replacement over the course of the 24-h and use the same 6 time points for the simulated pseudobulk datasets. For each “gene”, we randomly select 20% of the “single cells” without replacement and sum their expression to obtain a vector of pseudobulk replicates for that gene over the 6 selected time points. We repeat this procedure 500 times for each group, generating 500 pseudobulk datasets per group.

### 4.4 scRNA-seq experiments

#### 4.4.1 Mouse strains, husbandry

Wild type male C57BL/6 were housed under 12:12 light-dark cycle for two weeks prior to and during the time of experiment. To collect telogen skin, mice were about 54 days old by the time of sample collection.

#### 4.4.2 Sample collection and sequencing

Immediately after sacrificing a mouse with CO_2_, hair on dorsal skin was removed with an electric razor and Nair Hair Removal cream. After the dorsal skin is isolated from the body, fat and remaining blood vessels were scrapped away. A circular piece of skin was obtained with a 12mm biopsy punch, and minced into tiny pieces. 1x collagenase was then added to the minced skin and the suspension was incubated at 37 °C for 1.5 hours. The suspension is then filtered with a 70*μm* and a 40*μm* cell strainers to obtain single cells. SYTOX blue viability dye was then added to the cell suspension and live cells were sorted out using FACS at the UCI Stem Cell Research Center.

Samples were collected every four hours for three days to generate in total 18 sample. The Chromium Single Cell 3’ v3 (10x Genomics) libraries were prepared and sequenced by University of California Irvine Genomic High Throughput Facility with Illumina NovaSeq6000.

### 4.5 Datasets and analysis

#### 4.5.1 Preprocessing for benchmarking

For GSE56931, we subset the dataset provided in the TimeSignatR [30] package to only include the 24-hour normal baseline time points and filtered out the 38 hours of continuous wakefulness and subsequent recovery sleep. For GSE38622, expression matrix was normalized as explained in [33]. For GSE157077, we used the transcriptomes of the mice who were fed normal chow through an entire circadian cycle (24 hours). Since the mice were maintained on a 12-hour light/12-hour dark cycle, we chose to concatenate the three replicates of 24 hours each to create one set of samples over 72 hours. For GSE54650, the raw CEL files for the kidney and liver were imported using the function `read.celfiles` in R package oligo. Each raw data matrix was then normalized with Robust Multiarray Average (RMA) using the function `rma`. To map the GPL6246 platform ID REF to Ensembl transcript IDs, we used the transcript cluster ID and gene assignments listed in the table provided at https://www.ncbi.nlm.nih.gov/geo/query/acc.cgi?acc=GPL6246.

For each transcript cluster ID, we removed all gene assignments unless they were Ensembl transcript IDs or started with Gm. If a transcript cluster ID was mapped to more than one gene, then we replicated that row by the number of genes (e.g., transcript reference ID 10344614 is assigned to three Ensembl transcript IDs so that row in the normalized dataset was replicated three times). The expression for each transcript cluster ID was then divided by the number of genes assigned (e.g., since transcript reference ID 10344614 has three gene assignments, the values for all three rows in the normalized dataset were divided by three). Transcript cluster IDs that were not assigned to any genes were removed from the normalized dataset. To convert the Ensembl transcript IDs to gene names, we use R package biomaRt [58, 59]. If biomaRt did not find a gene name, then we kept the original Ensembl transcript ID. For GSE54651, we converted the Ensembl gene IDs to gene names using R package biomaRt [58, 59]. If biomaRt did not find a gene name, then we kept the original Ensembl gene ID. For each time point in GSE117295 and the scRNA-seq data we collected in this study, we summed the counts of each gene in all the cells without any pre-processing to create a pseudobulk dataset. In the case where the same gene occurs multiple times in the data, we took the mean of those entries. The resulting pseudobulk data at each time point is a single row vector in which each entry represents the expression value of a unique gene. The light-stimulated group is not considered in this paper.

#### 4.5.2 scRNA-seq data analysis for dermal skin

We used Cell Ranger version 3.1.0 with MM10 reference to process the raw sequencing output. The downstream analysis was done in Seurat V3 according to the vignette.

Cell with 850-7800 features and less than 13% of mitochondrial genes were kept. The SCTransform function was performed on each sample and 3250 integration features were selected using SelectIntegrationFeatures for each sample. Principal component analysis was then done on the integrated dataset and the Louvian algorithm was used to generate the clusters. Cluster identities were then determined in combination of marker genes found in the current clustering outcome and feature plots of canonical marker genes.

#### 4.5.3 Pseudobulk data analysis for dermal skin

meta2d from the MetaCycle package was used on the pseudobulk data generated from the scRNA-seq data collected from dermal skin to identify rhythmic genes. Genes with JTK pvalue*<*0.05 were determined to be significantly rhythmic. We used the meta2d phase column to split the rhythmic genes into four groups based on their peaking time.

Gene Ontology analysis was performed using ClusterProfiler in R with *p*-value *<* 0.05 as the significance cutoff.

### 4.6 Enrichment for GWAS SNPs

For rhythmic genes with JTK pvalue*<*0.01 in the dermal fibroblasts or immune cells, we used hypergeometric test to assess their enrichment among genes that are within 200kb of the GWAS signals of different skin immune-mediated conditions [43, 44, 60–64]. False discovery rate≤0.05 and observed-to-expected ratio≥2 are designated to be significant.

### 4.7 Statistics for circular data

The “circular” package was used to perform statistical calculations and tests, including calculation of the mean and standard deviation, as well as the Rao’s Tests for Homogeneity and the Wallraff Test of Angular Distances, for the circadian time prediction output in Section 2.7.

## 5 Data availability

All published datasets used in this paper can be accessed through their respective GEO accession codes. Although the datasets in [32] and [30] are both accessible through their GEO accession codes, this paper used the versions provided in the TimeSignatR package [30]: https://github.com/braunr/TimeSignatR.

The time series of scRNA-seq data from mouse dermal skin are available in the GEO database under GSE223109.

## 6 Code availability

tauFisher is available as an R package, which is available at https://github.com/micnngo/tauFisher (note that this will be a private repository until publication). The two methods we compared tauFisher against are also available as R packages: TimeSignatR at https://github.com/braunr/TimeSignatR and ZeitZeiger at https://github.com/hugheylab/zeitzeiger. For TimeSignatR, we modified the vignette the authors provided so that the two point calibration only calibrates the training dataset; the original vignette calibrates the concatenated training and test datasets.

## 7. Acknowledgements

This project is supported by an award entitled “Skin Biology Resource-Based Center at UCI Systems Biology Core” from the Center of Complex Biological Systems funded by NIH/NIAMS P30-AR075047 (J.D. and M.N.); National Institute of Health grants R01AR056439 and P30AR075047 (B.A.); NSF grant DMS1763272 and a grant from the Simons Foundation (594598) (J.D., M.N. and B.A.); National Institute of General Medical Sciences, National Research Service Award GM136624 (J.D.); NSF grant DMS1936833 (M.N. and B.S.); the California Institute for Regenerative Medicine Training Program Award EDUC4-12822 (S.S.K.); and Chan Zuckerberg Initiative grant DAF2022-239946 (B.A., J.G., and L.C.T.).

## 8 Author contributions

J.D., M.N., B.S., J.L. and B.A. conceived the project; J.D. and M.N. developed and implemented the tauFisher pipeline, and conducted the computational analysis; J.D. and S.S.K. collected the scRNA-seq data. L.T. and J.G. contributed to the GWAS SNPs enrichment analysis. J.D. and M.N. wrote the manuscript. All authors edited and approved the manuscript.

