## Supplemental Figure 1, 2; Supplemental Table 1, 2. for "tauFisher accurately predicts circadian time from a single sample of bulk and single-cell transcriptomic data"

### 1 Benchmark Training

Since tauFisher relies on the `meta2d` function in the MetaCycle R package [1] to identify the periodic genes, if a training set does not include consecutive samples, we modify the input training data to have `NA`s for those samples. This may occur when the training set includes time points such as $\{4, 12, 16, \cdots \}$ and the "missing" value, 8, is assigned to the testing set. We then insert a column for the "missing" value, 8, in the input data set for tauFisher and assign `NA`s to all genes in that column. For the kidney and liver bulk sequencing data sets [2], the `meta2d` function identifies a large number ($> 20$) of significant genes with a period of 24 hours. To reduce the number of genes, we apply the following filters in order: (1) filter out any genes that end with "Rik" or "-ps" or begin with "MT-" or "Gm", (2) has an amplitude greater than the mean amplitude, and (3) has an amplitude greater than the median amplitude.

### 2 Benchmark Results with Different Accuracy

**Table 1** Benchmark results (mean ± standard deviation) when we train on 80% of the data set and predict the circadian time of 20%.

| Data | Metric | ZeitZeiger | TimeSignature | | tauFisher | |
|---|---|---|---|---|---|---|
| | | | w/ 1-point | w/ 2-point | w/ Lomb-Scargle | w/ JTK_Cycle |
| [3] | Accuracy (within 3 hr) | 0.480 ± 0.122 | 0.250 ± 0.131 | 0.181 ± 0.098 | 0.456 ± 0.151 | **0.517 ± 0.135** |
| | Accuracy (within 2 hr) | 0.325 ± 0.103 | 0.168 ± 0.110 | 0.126 ± 0.084 | 0.347 ± 0.139 | **0.417 ± 0.148** |
| | Accuracy (within 1 hr) | 0.174 ± 0.094 | 0.085 ± 0.095 | 0.077 ± 0.076 | 0.209 ± 0.107 | **0.264 ± 0.129** |
| | Accuracy (exact) | 0.002 ± 0.011 | 0.000 ± 0.000 | 0.000 ± 0.000 | 0.076 ± 0.075 | **0.104 ± 0.078** |
| [4] | Accuracy (within 3 hr) | **0.635 ± 0.087** | 0.342 ± 0.097 | 0.266 ± 0.085 | 0.477 ± 0.096 | 0.625 ± 0.078 |
| | Accuracy (within 2 hr) | 0.487 ± 0.094 | 0.235 ± 0.087 | 0.181 ± 0.077 | 0.367 ± 0.088 | **0.499 ± 0.084** |
| | Accuracy (within 1 hr) | 0.250 ± 0.067 | 0.120 ± 0.061 | 0.084 ± 0.053 | 0.235 ± 0.073 | **0.304 ± 0.071** |
| | Accuracy (exact) | 0.001 ± 0.006 | 0.000 ± 0.000 | 0.000 ± 0.000 | 0.072 ± 0.053 | **0.092 ± 0.052** |
| [5] | Accuracy (within 3 hr) | 0.563 ± 0.350 | 0.320 ± 0.259 | 0.070 ± 0.136 | **0.783 ± 0.248** | 0.750 ± 0.286 |
| | Accuracy (within 2 hr) | 0.460 ± 0.344 | 0.210 ± 0.240 | 0.047 ± 0.116 | **0.740 ± 0.258** | 0.670 ± 0.330 |
| | Accuracy (within 1 hr) | 0.307 ± 0.275 | 0.107 ± 0.163 | 0.027 ± 0.091 | **0.590 ± 0.280** | 0.543 ± 0.295 |
| | Accuracy (exact) | 0.010 ± 0.057 | 0.000 ± 0.000 | 0.000 ± 0.000 | **0.257 ± 0.263** | 0.227 ± 0.241 |
| [6] | Accuracy (within 3 hr) | 0.108 ± 0.298 | 0.402 ± 0.230 | 0.100 ± 0.133 | 0.610 ± 0.237 | **0.705 ± 0.237** |
| | Accuracy (within 2 hr) | 0.075 ± 0.218 | 0.285 ± 0.228 | 0.075 ± 0.120 | 0.545 ± 0.239 | **0.635 ± 0.237** |
| | Accuracy (within 1 hr) | 0.040 ± 0.150 | 0.128 ± 0.172 | 0.032 ± 0.084 | 0.412 ± 0.257 | **0.468 ± 0.253** |
| | Accuracy (exact) | 0.000 ± 0.000 | 0.000 ± 0.000 | 0.000 ± 0.000 | **0.190 ± 0.195** | 0.182 ± 0.181 |
| [2][2,4] | Accuracy (within 3 hr) | 0.996 ± 0.028 | 0.256 ± 0.178 | 0.192 ± 0.133 | 0.996 ± 0.028 | **1.000 ± 0.000** |
| | Accuracy (within 2 hr) | 0.986 ± 0.051 | 0.176 ± 0.159 | 0.100 ± 0.119 | 0.966 ± 0.086 | **1.000 ± 0.000** |
| | Accuracy (within 1 hr) | 0.728 ± 0.179 | 0.096 ± 0.125 | 0.012 ± 0.048 | 0.880 ± 0.158 | **0.916 ± 0.128** |
| | Accuracy (exact) | 0.004 ± 0.028 | 0.000 ± 0.000 | 0.000 ± 0.000 | **0.370 ± 0.219** | 0.358 ± 0.209 |
| [2][1,2,5] | Accuracy (within 3 hr) | NA | 0.475 ± 0.279 | 0.115 ± 0.211 | 0.720 ± 0.416 | **0.945 ± 0.200** |
| | Accuracy (within 2 hr) | NA | 0.330 ± 0.286 | 0.100 ± 0.201 | 0.695 ± 0.414 | **0.945 ± 0.200** |
| | Accuracy (within 1 hr) | NA | 0.240 ± 0.251 | 0.070 ± 0.174 | 0.565 ± 0.453 | **0.815 ± 0.346** |
| | Accuracy (exact) | NA | 0.000 ± 0.000 | 0.000 ± 0.000 | 0.250 ± 0.314 | **0.375 ± 0.344** |
| [2][3,4] | Accuracy (within 3 hr) | 0.964 ± 0.100 | 0.264 ± 0.186 | 0.168 ± 0.105 | 0.936 ± 0.133 | **0.986 ± 0.059** |
| | Accuracy (within 2 hr) | 0.868 ± 0.156 | 0.164 ± 0.137 | 0.076 ± 0.106 | 0.880 ± 0.161 | **0.932 ± 0.103** |
| | Accuracy (within 1 hr) | 0.626 ± 0.225 | 0.086 ± 0.118 | 0.000 ± 0.000 | 0.764 ± 0.196 | **0.850 ± 0.162** |
| | Accuracy (exact) | 0.002 ± 0.020 | 0.000 ± 0.000 | 0.000 ± 0.000 | **0.312 ± 0.189** | 0.308 ± 0.219 |
| [2][1,3,5] | Accuracy (within 3 hr) | NA | 0.465 ± 0.357 | 0.075 ± 0.179 | 0.840 ± 0.332 | **0.970 ± 0.171** |
| | Accuracy (within 2 hr) | NA | 0.360 ± 0.341 | 0.065 ± 0.169 | 0.765 ± 0.344 | **0.910 ± 0.269** |
| | Accuracy (within 1 hr) | NA | 0.240 ± 0.289 | 0.060 ± 0.163 | 0.585 ± 0.427 | **0.735 ± 0.313** |
| | Accuracy (exact) | NA | 0.000 ± 0.000 | 0.000 ± 0.000 | 0.155 ± 0.263 | **0.335 ± 0.326** |

[1] If ZeitZeiger [7] was unable to do a 3-fold cross validation, we ran ZeitZeiger without any cross validation and set `sumabsv` = 1 and `nSpc` = 3.

[2] kidney

[3] liver

[4] microarray

[5] bulk RNA

### 3 Benchmark Result Table

**Table 2** Benchmark results (mean ± standard deviation) when we train on 80% of the data set and predict the circadian time of 20%.

| Data | Metric | ZeitZeiger | TimeSignature | | tauFisher | |
|---|---|---|---|---|---|---|
| | | | w/ 1-point | w/ 2-point | w/ Lomb-Scargle | w/ JTK_Cycle |
| [3] | Accuracy | 0.325 ± 0.103 | 0.168 ± 0.110 | 0.126 ± 0.084 | 0.347 ± 0.139 | **0.417 ± 0.148** |
| | RMSE | **4.829 ± 0.779** | 7.213 ± 1.440 | 7.209 ± 1.403 | 5.666 ± 0.848 | 5.167 ± 0.890 |
| | # NA | 0 | 6 | 6 | 0 | 0 |
| [4] | Accuracy | 0.487 ± 0.094 | 0.235 ± 0.087 | 0.181 ± 0.077 | 0.367 ± 0.088 | **0.499 ± 0.084** |
| | RMSE | **3.957 ± 0.571** | 6.198 ± 0.574 | 6.572 ± 0.517 | 5.606 ± 0.646 | 4.655 ± 0.610 |
| | # NA | 0 | 0 | 0 | 0 | 0 |
| [5] | Accuracy | 0.460 ± 0.344 | 0.210 ± 0.240 | 0.047 ± 0.116 | **0.740 ± 0.258** | 0.670 ± 0.330 |
| | RMSE | 4.558 ± 3.640 | 5.781 ± 1.785 | 7.982 ± 1.949 | **2.488 ± 1.662** | 2.707 ± 1.976 |
| | # NA | 15 | 0 | 0 | 0 | 0 |
| [6] | Accuracy | 0.075 ± 0.218 | 0.285 ± 0.228 | 0.075 ± 0.120 | 0.545 ± 0.239 | **0.635 ± 0.237** |
| | RMSE | 10.784 ± 3.317 | 5.540 ± 1.512 | 7.883 ± 1.392 | 3.722 ± 1.880 | **2.969 ± 1.477** |
| | # NA | 88 | 0 | 0 | 0 | 0 |
| [2][2,4] | Accuracy | 0.986 ± 0.051 | 0.176 ± 0.159 | 0.100 ± 0.119 | 0.966 ± 0.086 | **1.000 ± 0.000** |
| | RMSE | **0.849 ± 0.244** | 6.639 ± 1.366 | 6.511 ± 1.172 | 1.021 ± 0.406 | 0.911 ± 0.256 |
| | # NA | 0 | 0 | 0 | 0 | 0 |
| [2][1,2,5] | Accuracy | NA | 0.330 ± 0.286 | 0.100 ± 0.201 | 0.695 ± 0.414 | **0.945 ± 0.200** |
| | RMSE | NA | 4.578 ± 1.733 | 8.136 ± 2.638 | 2.965 ± 3.117 | **1.076 ± 0.981** |
| | # NA | 100 | 0 | 0 | 0 | 0 |
| [2][3,4] | Accuracy | 0.868 ± 0.156 | 0.164 ± 0.137 | 0.076 ± 0.106 | 0.880 ± 0.161 | **0.932 ± 0.103** |
| | RMSE | 1.391 ± 1.124 | 6.467 ± 1.382 | 6.844 ± 1.137 | 1.797 ± 1.479 | **1.182 ± 0.451** |
| | # NA | 0 | 0 | 0 | 0 | 0 |
| [2][1,3,5] | Accuracy | NA | 0.360 ± 0.341 | 0.065 ± 0.169 | 0.765 ± 0.344 | **0.910 ± 0.269** |
| | RMSE | NA | 3.787 ± 2.067 | 8.343 ± 2.434 | 2.269 ± 2.356 | **1.203 ± 0.825** |
| | # NA | 100 | 0 | 0 | 0 | 0 |

[1]If ZeitZeiger [7] was unable to do a 3-fold cross validation, we ran ZeitZeiger without any cross validation and set `sumabsv` = 1 and `nSpc` = 3.

[2]kidney

[3]liver

[4]microarray

[5]bulk RNA

### 4 Supplementary figures

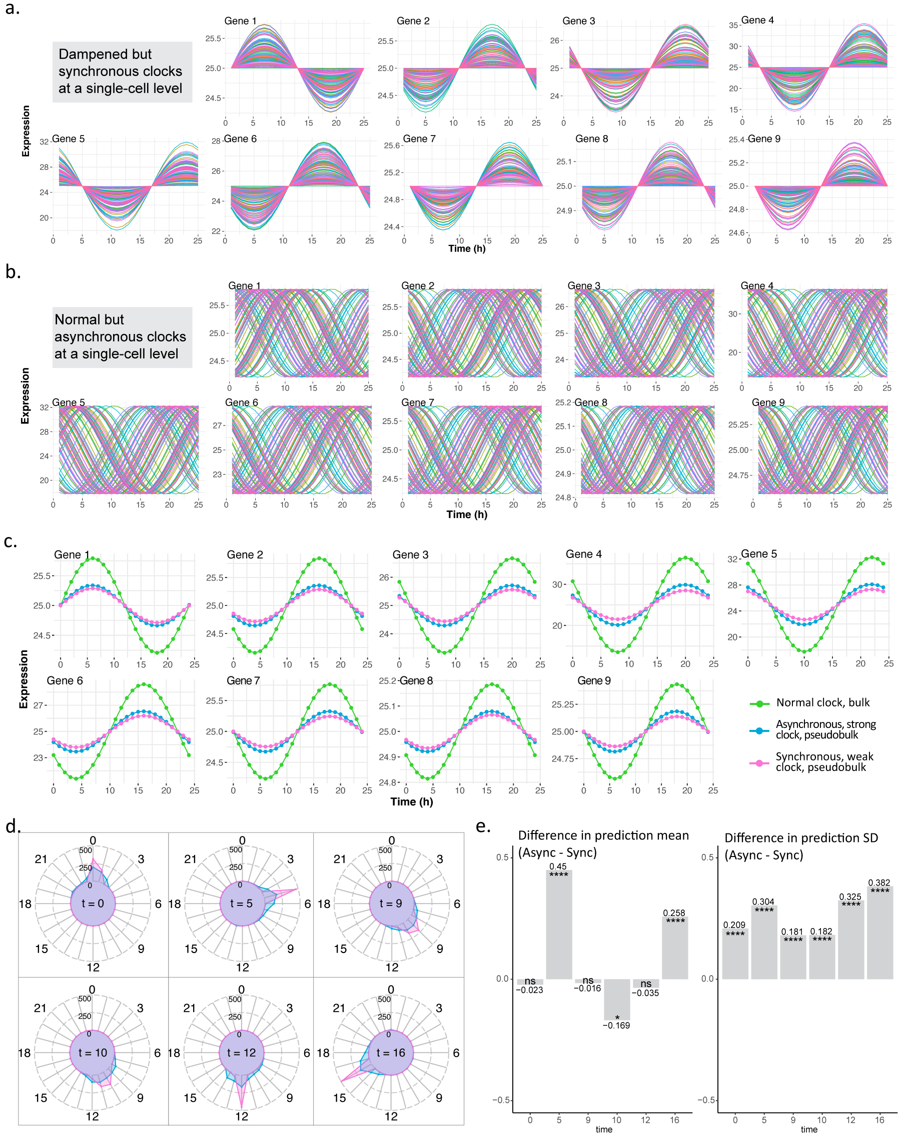


**Fig. 1** We tested tauFisher's ability to determine circadian phase heterogeneity on simulated data. **a** Simulated expression of nine predictor genes in 100 single cells that have synchronous circadian clocks but dampened amplitudes. Each line represents gene expression a cell over time. **b** Simulated expression of nine predictor genes in 100 single cells that have asynchronous circadian clocks but regular amplitudes. Each line represents gene expression a cell over time. **c** At the bulk level, scenarios in **a** and **b** generate similar patterns, which are oscillations with dampened amplitudes but similar peaking time when compared to a group of cells with normal clocks. **d** We combined tauFisher with bootstrapping and randomly selected six time points to do the comparison. We generated 500 time predictions and plotted the distributions for the cells in **a** (red) and **b**(blue). **e** Bar plots showing the differences between the prediction mean (left) and standard deviation (right) at each time point (cells in **b** - cells in **a**). ns: $p$-value $> 0.05$, $*$: $p$-value $\leq 0.05$, $p$-value $\leq 0.01$, $***$: $p$-value $\leq 0.001$, $****$: $p$-value $\leq 0.0001$.

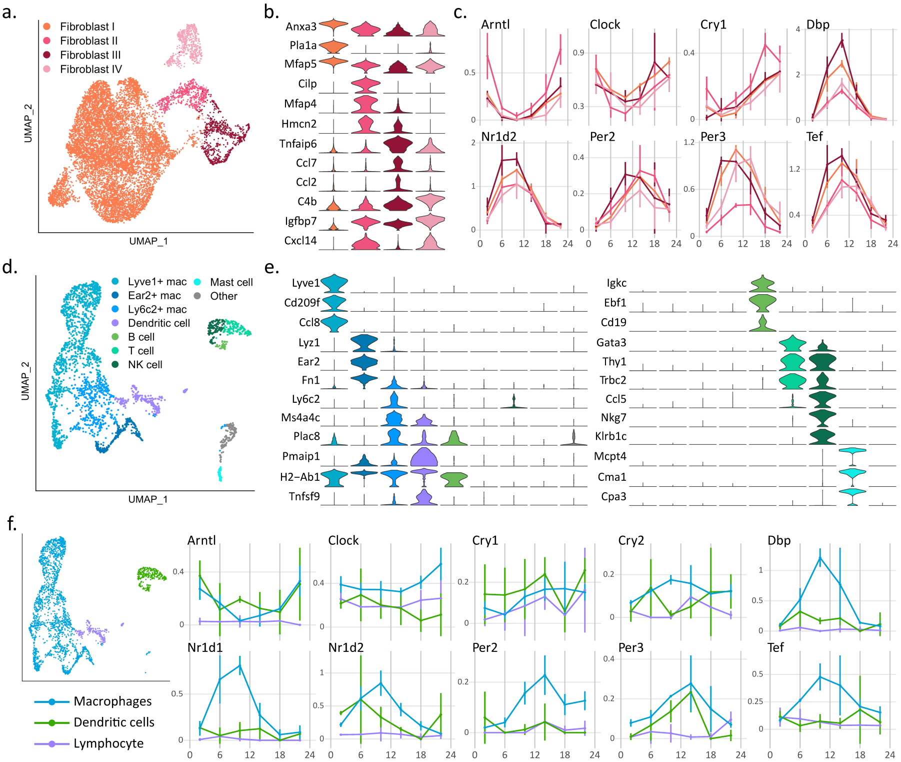


**Fig. 2** Subclustering results for the mouse dermal fibroblasts and immune cells. **a** Four subclusters were identified for the dermal fibroblasts. **b** Violin plots show expression of marker genes for each cluster. **c** Core clock gene expression for the four fibroblast subclusters show similar rhythmic pattern. **d** Eight subclusters including both myeloid cells and lymphoid cells. **e** Violin plots show expression of marker genes for each cluster. **f** Due to the low cell count for some time points, we combined the eight subclusters of immune cells form three major clusters. The core clock gene expression over time is plotted for the three major clusters. With the great variability present in the data, probably contributed by low cell counts, the core clock genes do not show robust rhythms.
